# CSF1R inhibition by a small molecule inhibitor affects hematopoiesis and the function of macrophages

**DOI:** 10.1101/2019.12.27.889469

**Authors:** Fengyang Lei, Naiwen Cui, Chengxin Zhou, James Chodosh, Demetrios G. Vavvas, Eleftherios I. Paschalis

**Affiliations:** Massachusetts Eye and Ear, Department of Ophthalmology, Harvard Medical School, Boston, MA; Boston Keratoprosthesis Laboratory, Harvard Medical School, Boston, MA; Disruptive Technology Laboratory, Massachusetts Eye and Ear, Department of Ophthalmology, Harvard Medical School, Boston, MA; John A. Paulson School of Engineering and Applied Sciences, Harvard University, Cambridge, MA 02138, USA; Angiogenesis Laboratory, Massachusetts Eye and Ear, Department of Ophthalmology, Harvard Medical School, Boston, MA

**Keywords:** Eye, neuroglia, microglia, macrophage, remodeling, CSF1R

## Abstract

Colony-stimulating factor 1 receptor (CSF1R) inhibition has been proposed as a method for microglia depletion, with the assumption that it does not affect peripheral immune cells. Here, we show that CSF1R inhibition by PLX5622 indeed affects the myeloid and lymphoid compartments, causes long-term changes in bone marrow-derived macrophages by suppressing their IL-1β, CD68 and phagocytosis, but not CD208, following exposure to endotoxin, and also reduces the population of tissue resident macrophages of peritoneum, lung, liver, but not spleen. Thus, small molecule CSF1R inhibition is not restricted to microglia only, but rather causes strong effects on circulating and tissue macrophages that perdure long after cessation of the treatment. Given that peripheral monocytes repopulate the CNS after CSF1R inhibition, these changes may have practical implications on relevant experimental data.

---

Colony stimulating factor 1 receptor (CSF1R) inhibition has been proposed as a specific method for microglia that does not affect peripheral immune cells(1-4). However, this claim has been based solely on cell count measurements of blood monocytes and evaluation of blood brain barrier rather than direct assessment of cellular subtypes and their function. Also, recent data have shown an effect on liver and lung tissue resident macrophage (5, 6) Given that peripheral monocytes have been shown to participate in CNS disease via both infiltration and repopulation of neuroglia following microglia depletion(4, 7-12), it is important to determine if CSF1R inhibition causes functional changes in peripheral immune cells that can become part of the CNS(3, 4, 12-16).

Here we show that, contrary to the accepted notion(1, 13), PLX5622, a commonly used small molecule CSF1R inhibitor(10, 12, 13), does not affect only microglia but also leads to long-term changes in the myeloid and lymphoid compartments of the bone marrow, spleen and blood by suppressing CCR2^+^ monocyte progenitor cells, CX3CR1^+^ bone marrow-derived macrophages (BMDM), CD117^+^ (C-KIT^+)^ hematopoietic progenitor cells, F4/80^+^, MerTK^+^ and CD34^+^ hematopoietic stem cells **(fig. 1 A-G)**. Most importantly, these cell populations either do not recover or rebound after cessation of CSF1R inhibition, with the exception of CD45^+^ CD11b^+^ cells which remain unaffected **(fig. 1 A-G)**. Besides the effects on the myeloid compartment, CSF1R inhibition also alters the lymphoid compartment of the bone marrow by suppressing T-cells (CD3^+^, CD4^+^, and CD8^+^), **(fig. 1 G)** and by upregulating CD19^+^ B cells **(fig. 1 G)**. Cessation of CSF1R inhibition causes rebound of some, but not all, lymphoid cells (**fig. 1 G)**.

**Fig. 1.**
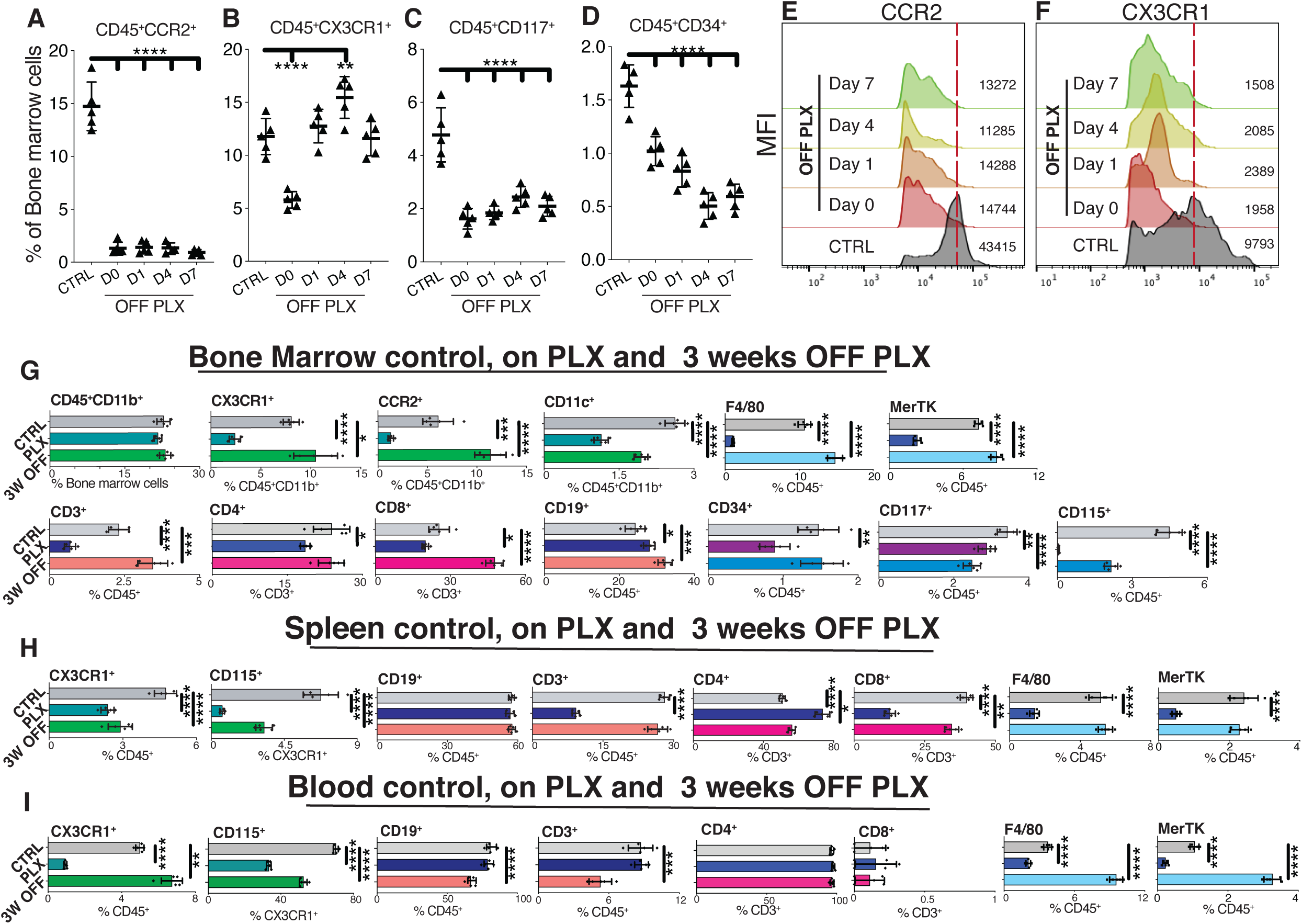
CSF1R inhibition by PLX5622 affects the myeloid and lymphoid compartments of the bone marrow, spleen and blood. Flow cytometric analysis of bone marrow cells isolated from CCR2^+/RFP^::CX3CR1^+/GFP^ mice immediately, after 3-week treatment with CSF1R inhibitor (PLX5622), and at different timepoints after cessation of the CSF1R inhibitor. **(A - F)** CSF1R inhibition suppresses CCR2^+^, CX3CR1^+^, CD117^+^, and CD34^+^ cells. One week after cessation of CSF1R inhibitor, only macrophages recover in number, although with a lower expression of CX3CR1. **(G)** CSF1R inhibition does not affect CD45^+^, CD11b^+^ and Ly6C^+^ bone marrow myeloid cell populations, but does suppress CD11c^+^ dendritic cells, CD4^+^ and CD8^+^ T lymphocytes, as well as, CD115^+^, CD117^+^ and CD34^+^ hematopoietic subsets, and upregulates CD19^+^ B cells. Three weeks after cessation of CSF1R inhibition, CX3CR1^+^, CCR2^+^, Ly6C^+^ CD3^+^ and CD8^+^ sub-populations rebound; Ly6G^+^ granulocytes, CD115^+^, and CD117^+^ cells remain suppressed; CD4^+^ T cells and CD34^+^ cells recover; and CD19^+^ B cells remain upregulated. **(H)** Effects of CSF1R inhibition on the spleen’s myeloid and lymphoid populations. Only CD19^+^ B cells remain unaffected. **(I)** CSF1R inhibition causes immediate suppression in the myeloid compartment late suppression of the lymphoid compartment of the blood. n=5 per group, mean±SD, One-way analysis of variance with Dunnett’s correction for multiple comparisons, *\* P<0.05, ** P<0.01, *** P<0.001, **** P<0.0001*.

In addition to the effects on bone marrow cells, CSF1R inhibition also suppresses splenic CX3CR1^+^ cells and this inhibition persist for at least 3 weeks after cessation of treatment **(fig. 1 H)**. Moreover, splenic CD3^+^ T cells (primarily CD8^+^) become suppressed, whereas CD19^+^ B cells are not affected by the inhibitor **(fig. 1 H)**.

Likewise, CSF1R inhibition suppresses circulating CX3CR1^+^, CD115^+^, F4/80^+^ and MerTK^+^ blood cells with no immediate effect on the lymphoid CD3^+^ and CD4^+^ populations **(fig. 1 I)**. Cessation of CSF1R inhibitor causes rebound of the CX3CR1^+^, F4/80^+^ and MerTK^+^, but not CD115^+^ blood cells, and leads to delayed suppression of CD19^+^, CD3^+^ and CD4^+^ lymphoid cell in the circulation **(fig. 1 I)**.

CSF1R inhibition also suppresses the proliferation of bone marrow and spleen macrophages **(fig. 2 A-D)**, and impairs the function of bone marrow-derived macrophages (BMDM) for the long-term. In fact, 3 weeks after cessation of CSF1R inhibition, BMDMs display reduced IL-1β expression in response to endotoxin **(fig. 2 E, F)**, diminished phagocytosis **(fig. 2 E-G)**, reduced CD68, but not CD206 expression **(fig. 2 I-L)**, and suppressed CD115 labeling **(fig. 2 M-N)**. Moreover, CSF1R inhibition reduces the number of tissue resident CX3CR1^+^ femur, CD11a^+^ lung, CD102^+^ peritoneum, MHC-II^hi/lo^ liver, but not CD106^+^ spleen macrophages, while consistently suppresses bone marrow-derived CX3CR1^+/EGFP^ macrophages in these tissues **(fig. 2 O)**.

**Fig. 2.**
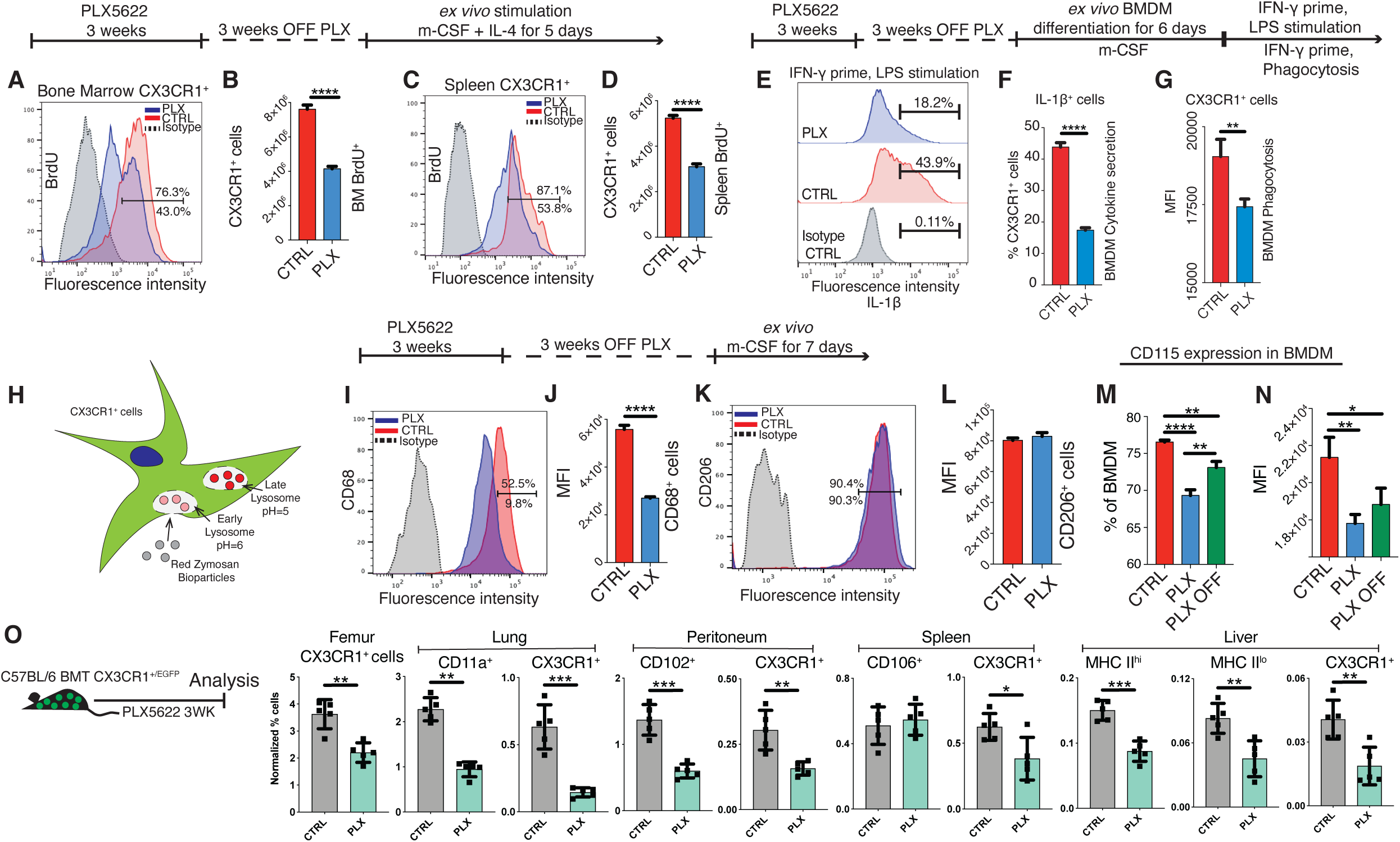
CSF1R inhibition by PLX5622 affects the function and survival of tissue resident macrophages of the bone marrow, lung, peritoneum, spleen and liver. **(A-D)** *ex vivo* evaluation of the function of bone marrow-derived macrophage (BMDM) from CX3CR1^+/GFP^ mice 3 weeks after cessation of CSF1R inhibitor. Macrophages from the bone marrow or spleen exhibit reduced proliferation 3 weeks after cessation of CSF1R inhibition. **(E-L)** CSF1R inhibition suppresses IL-1β, CD68 expression and phagocytosis of bone marrow-derived macrophage following exposure to lipopolysaccharide, but does not affect their CD206 expression. **(H)** Schematic representation of the phagocytosis assay. **(M, N)** CSF1R inhibition causes long-term suppression of CD115 macrophage marker. **(O)** CSF1R inhibition reduces the number of tissue resident CX3CR1^+^ femur, CD11a^+^ lung, CD102^+^ peritoneum, MHC-II^hi/lo^ liver, but not CD106^+^ spleen macrophages, while consistently suppresses bone marrow-derived CX3CR1^+/EGFP^ macrophages in these tissues. n=5 per group, mean±SD, Independent t-test, *\*P<0.05, ** P<0.01, ***P<0.001, **** P<0.0001*.

Previous studies have suggested that microglia depletion by CSF1R inhibition can either promote or exacerbate neurodegeneration(4, 12, 17). This puzzling and contradictory results may not only be due to the differing role of microglia in various disease models but also due to the varying relative contribution of peripheral and circulating macrophages on disease phenotype. Our study demonstrates that CSF1R inhibition affects both the circulating and tissue resident macrophages of various tissue which may explain in part the reason for the discrepancy in recent investigations.

Considering that BMDMs permanently engraft into diseased CNS tissue(10, 12), this work suggests that small molecule inhibition is not restricted to microglia but additionally affects the turnover and function of bone marrow-derived, circulating and tissue resident macrophages. These effects perdure long after cessation of the treatment and has implications in the interpretation of relevant experimental data.

## Materials and Methods

### Mouse model

All animal experiments were performed in accordance with the Association for Research in Vision and Ophthalmology Statement for the Use of Animals in Ophthalmic and Vision Research, and the National Institutes of Health (NIH) Guidance for the Care and Use of Laboratory Animals. This study was approved by the Mass. Eye and Ear Animal Care Committee. Mice at 6-12 months old were used: C57BL/6J (Stock#: 000664), B6.129(Cg)-Ccr2tm2.1lfc/J (Stock#: 017586) and B6.129P-Cx3cr1tm1Litt/J (Stock#: 005582) Jackson Laboratory. CCR2^RFP/+^::CX3CR1^EGFP/+^ generated by crossing B6.129(Cg)-Ccr2tm2.1lfc/J with B6.129P-Cx3cr1tm1Litt/J. CX3CR1^EGFP/+^ generated by crossing male B6.129P-Cx3cr1tm1Litt/J with female C57BL/6J. Mice were bred in house. Microglia depletion was performed by chow administration for 3 weeks of PLX5622 (Plexxikon Inc., Berkeley, CA). Flow cytometry, and *ex vivo* BMDM evaluation was performed as previously described(10, 12). Blood cells was collected by cardiac puncture and centrifugation. Peritoneum, lung, liver, spleen and femur resident macrophage were evaluated using appropriate markers (18, 19) in BMT CX3CR1^+/EGFP^ reporter mice following 3-weeks exposure to PLX5622.

### Flow cytometry markers

Bone marrow and spleen cells from CX3CR1^+/GFP^ and CX3CR1^+/EGFP^::CCR2^+/RFP^ reporter mice were blocked with CD16/32 (Clone: 2.4G2), analyzed with IL-1β (Clone: NJTEN3), Lyve1 (Clone: ALY7) eBiosciences (San Diego, CA); CD45 (Clone: 104), CD11b (Clone: M1/70), CD11c (Clone: N418), CD3 (Clone: 17A2), CD4 (Clone: GK1.5), CD8 (Clone: 53-5.8), CD19 (Clone: 6D5), CD117 (Clone:2B8), CD34 (Clone: HM34), CD115 (Clone: AFS98), CD68 (Clone: FA-11), CD206 (Clone: C068C2), CCR2(Clone: SA203G11), BrdU(Clone: Bu20a), F4/80 (Clone: BM8), MerTK (Clone: 2B10C42), CD11a (Clone: I21/7), CD102 (Clone: 3C4), CD106 (Clone: 429), and I-A/I/E (Clone: M5/114.15.2) BioLegend (San Diego, CA). Intracellular staining was performed by fixing cells in Paraformaldehyde-based Fixation buffer (BioLegend) followed by permeabilization with Perm/Wash buffer (BioLegend). Cells were analyzed on a BD LSR II cytometer (BD Biosciences, San Jose, CA, USA) using FlowJo software (Tree Star, Ashland, OR, USA).

### LPS stimulation assay

BMDM were primed with 150U/mL interferon-γ (IFNγ) for 6 hours followed by LPS at final concentration of 10ng/mL (Sigma-Aldrich, St. Louis, MO) added into culture medium for 20 hours. Brefeldin A 5μg/mL (BD Pharmingen, Bedford, MA) was added 4 hours before cell harvest and flow cytometry.

### Phagocytosis Assay

The pHrodo^™^ Red BioParticles® Conjugates for Phargocytosis (P35364) (Molecular Probes, Eugene, OR) kit was used. Six days after cell plating with mCSF, and one day prior to the assay, BMDMs were recovered from culture and seeded. Cells were stimulated with IFNγ for 4 hours and then culture medium was replaced with reconstituted red Zymosan A BioParticles. Cells were incubated at 37°C for 2 hours, trypsinized, and evaluated with flow cytometry.

### Statistical analysis

Data were analyzed with GraphPad (Prism 2.8.1, San Diego, CA) using two-tailed unpaired t-test and ordinary one-way ANOVA with Dunnet’s correction for multiple comparisons. Statistical significance was determined at P < 0.05.

## Author contributions

FL designed experiments, acquired data and analyzed data; NC, CZ analyzed data; DGV wrote and reviewed the manuscript; JC reviewed the manuscript; EIP designed experiments, analyzed data, and wrote the manuscript.

## Acknowledgments

This work was supported by the Boston Keratoprosthesis Research Fund, Massachusetts Eye and Ear, the Eleanor and Miles Shore Fund, the Massachusetts Lions Eye Research Fund, an unrestricted grant to the Department of Ophthalmology, Harvard Medical School, from Research to Prevent Blindness, NY, NY, NIH National Eye Institute core grant P30EY003790; Yeatts Family Foundation; Monte J Wallace Chair; Macula Society Research Grant award; an RPB Physician Scientist Award; NEI R21EY023079; NEI grant EY014104 (MEEI Core Grant); Loeffler Family fund; R01EY025362-01; ARI Young investigator Award. PLX5622 was kindly provided by Plexxikon Inc.

